# Monocular deprivation during the critical period alters neuronal tuning and the composition of visual circuitry

**DOI:** 10.1101/2022.08.23.504998

**Authors:** Thomas C. Brown, Aaron W. McGee

**Affiliations:** Department of Anatomical Sciences and Neurobiology, School of Medicine; University of Louisville, Louisville, KY, 40202

## Abstract

Abnormal visual experience during a developmental critical period degrades cortical responsiveness. Yet how experience-dependent plasticity alters the response properties of individual neurons and composition of visual circuitry is unclear. Here we measured with calcium imaging in alert mice how monocular deprivation (MD) during the developmental critical period affects binocularity, orientation, and spatial frequency tuning for neurons in primary visual cortex. Tracking the tuning properties for several hundred neurons revealed that the interconversion of monocular and binocular neurons relies on the quality of visual experience to determine the ratio of monocular neurons responsive to the contralateral and ipsilateral eye. In addition, a population of neurons more responsive to the closed eye were exchanged for neurons with tuning properties more similar to the responsive neurons altered by MD. Thus, plasticity during the critical period adapts to recent experience by both altering the tuning of responsive neurons and recruiting neurons with matching tuning properties.

## Introduction

The impact of experience on maturing neural circuitry is most profound during a ‘critical period’ of heightened sensitivity in development (Levelt and Hübener, 2012). How experience sculpts neural circuits has been studied extensively in the primary visual cortex (V1), where monocular deprivation (MD) during a critical period perturbs the response properties of neurons (Espinosa and Stryker, 2012; Hensch and Quinlan, 2018). The effects of MD on visual cortex were first characterized cats and primates as ‘single units’ by auditory discrimination (Hubel et al., 1977; Hubel and Wiesel, 1970; Wiesel and Hubel, 1963). Experiments of similar design were later performed with rodents (Drager, 1978; Fagiolini et al., 1994; Gordon and Stryker, 1996). Units were typically only examined for ocular dominance (OD) because quantifying additional tuning properties manually was arduous.

Subsequent studies in the rodents have employed techniques to study OD plasticity that report aggregate neuronal activity, such as visually-evoked potentials (VEPs), optical imaging of intrinsic signals, and multi-unit electrophysiologic recordings (Cang et al., 2005; Frenkel and Bear, 2004; Stephany et al., 2014). These approaches have elucidated several characteristics of OD plasticity during the critical period that discriminate it from the less robust plasticity remaining in the adult brain after the critical period closes (Morishita and Hensch, 2008). OD plasticity during the critical period is predominantly driven by an overall depression of responsiveness to the deprived eye (Frenkel and Bear, 2004; Sato and Stryker, 2008). Shifts in binocularity are also preceded by intracortical disinhibition and are insensitive to benzodiazepines and barbiturates. In contrast, OD plasticity in adults is smaller in magnitude, results from potentiation of responses to the fellow eye, and is not associated with disinhibition and is suppressed by benzodiazepines and barbiturates (Hensch et al., 1998; Kuhlman et al., 2013; Pham et al., 2004; Sato and Stryker, 2008; Sawtell et al., 2003; Stephany et al., 2016). However, these techniques are not suitable for measuring tuning properties at neuronal resolution do not inform how the tuning of individual neurons changes during OD plasticity.

Calcium imaging provided a first glimpse at how large populations of neurons in V1 responded to MD. Initial studies reported in layer (L) 2/3 of the binocular zone of mouse V1 are almost exclusively binocular (Mrsic-Flogel et al., 2007). However, more recent imaging experiments with either calcium-sensitive dyes or the genetically-encoded calcium sensor GCaMP6s have reported that a majority of neurons are monocular (Jenks and Shepherd, 2020; Salinas et al., 2017; Scholl et al., 2017; Tan et al., 2020). The orientation and spatial frequency (SF) tuning of excitatory neurons in V1 in juvenile and adult mice have also been characterized separately from both electrophysiology recordings sorted to isolate responses of individual neurons and calcium imaging. These studies measured the orientation tuning width and binocular matching for orientation, and tuning range and width for SF from naïve animals (Durand et al., 2016; Jeon et al., 2018; Marshel et al., 2011; Mrsic-Flogel et al., 2007; Niell and Stryker, 2008; Salinas et al., 2017; Scholl et al., 2017; Tan et al., 2020; Wang et al., 2010). Interestingly, brief MD during the critical period reduces orientation tuning in cats but not mice (Crair et al., 1997; Wang et al., 2010).

The effects of MD on binocularity and orientation tuning in adult mice has been tracked over weeks for a relatively small number of neurons per mouse with calcium imaging (Rose et al., 2016). This difficult repeat imaging study identified aspects of how MD alters the binocularity of population of neurons pooled across adult mice, as well as how these neurons recover as a population following restoration of binocular vision following MD. This study also measured the preferred orientation for a few dozen neurons and determined that it was not affected by MD.

More recent work has demonstrated a surprising reorganization of the neuronal composition of visual circuitry in mice throughout life as poorly tuned binocular neurons are rendered monocular and sharply tuned monocular neurons convert to binocular neurons (Tan et al., 2020). This process is disrupted by dark exposure during the critical period, a procedure that does not engage OD plasticity (Kang et al., 2013). Yet how abnormal vision during the critical period alters the interconversion of monocular and binocular neurons, the population of neurons active in visual circuitry, or neuronal tuning properties remains unknown. Here we investigated with calcium imaging how disrupting visual experience by MD during the critical period affects the neuronal composition of visual circuitry and neuronal tuning properties.

## Results

We implanted cranial windows at (postnatal (P) day 24-30) in mice expressing GCaMP6s in excitatory neurons of the forebrain (21, 22) (Figure S1A). The next day we identified the binocular zone of visual cortex by wide-field calcium imaging (22) (Figure S1B). Thereafter, we measured calcium responses in alert mice in response to a battery of sinusoidal gratings across a range of orientation (0-180 degrees, 30 degree intervals) and SF (.02 to .48 cycles per degree (cpd), intervals spaced at half octaves (log(1.5)) (Figure S1C,D). Visual stimuli were presented independently to each eye.

To measure the tuning properties in Layers (L) 2/3 and L4 in V1, first we segmented neuronal soma as regions of interest (ROIs) and then determined the z-score and inferred spike rate (ISR) for each ROI from the normalized change in fluorescence (dF/F) for each combination of orientation and SF (Figure S1E-I) (Berens et al., 2018; Ringach et al., 2016). We identified visually-responsive neurons from these ROIs based on the delay from stimulus onset (in imaging frames) and signal to noise ratio (SNR)(Tan et al., 2020), and well as the number of responses (spike ratio, SR) at the optimal delay (Figure S1J-N). The preferred orientation and SF were calculated from the row (orientation) and column (SF) corresponding to the maximal ISR at the optimal delay (frame number). In total, we evaluated calcium responses from 5557 neurons in this experiment, 2723 (~50%) of which met our inclusion criteria for visually-responsive neurons.

We calculated the ocular dominance index (ODI) for each neuron from dF/F for the visual stimulus capturing the preferred orientation and SF for non-deprived mice at P28 and P32, as well as mice at P32 after 4 days of MD (Figure 1A). Non-deprived mice were more responsive to visual stimuli presented to the contralateral eye, with higher average ODI values per mouse. By comparison, mice receiving 4-day MD displayed lower average ODI values (P< .0001) (Figure 1B). In addition, we calculated ODI score for each mouse from the sum of the dF/F values for the contralateral and ipsilateral eye from all visually responsive neurons (Figure 1C). This metric integrates the response strength for each neuron to evaluate OD. Contralateral bias was similarly reduced in mice following 4 days of MD (P < .0001). Thus, OD plasticity can be measured by calcium imaging either as the average ODI of individual neurons per mouse or the sum of dF/F for all responsive neurons per mouse.

**Figure 1.**
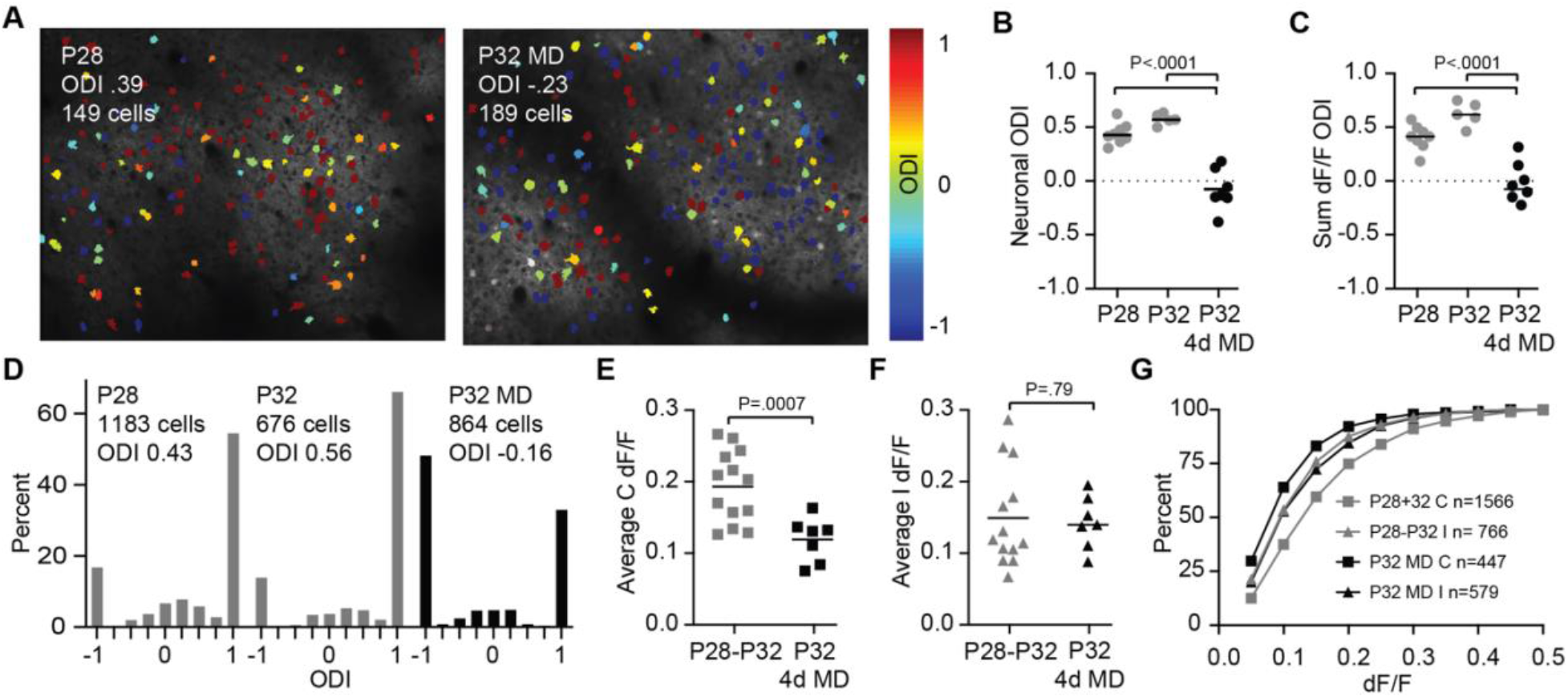
Measuring OD plasticity with calcium imaging at cellular resolution. (**A**) Imaging fields for P28 (*left*) and P32 after 4 days of MD (*right*). Neurons are color-coded to their Ocular Dominance Index (ODI) score. 1 corresponds to monocular contralateral neurons. −1 corresponds to monocular ipsilateral neurons. (**B**) Average ODI scores for non-deprived P28-P32 mice (mean ODI = 0.43, n= 8 mice), P32 mice (mean ODI .57, n= 5 mice), and P32 mice after 4 days of MD (mean ODI = −.09, n= 7 mice. P32 mice after 4 days of MD possess significantly different ODI values than both P28 and P32 non-deprived mice. (1-way ANOVA) **(B)** Sum ODI scores for the same mice as in panel B. P28 (mean ODI = 0.39), P32 (mean ODI=.61), and P32MD (mean ODI = −.02). P32 mice (mean ODI .57, n= 5 mice), and P32 mice after 4 days of MD (mean ODI = −.09, n= 7 mice. P32 mice after 4 days of MD possess significantly different ODI values than both P28 and P32 non-deprived mice. (1-way ANOVA). (**C**) Histogram of ODI scores for non-deprived P28 (*left*), non-deprived P32 mice (*middle*) and P32 mice after 4 days (*right*) P32 mice following 4 days (4d) of MD (*right*) from panel B. (**E, F**) Mean delta F over F (dF/F) values neurons responding to the contralateral eye (C) and ipsilateral eye (I) for each non-deprived mouse (P28-32, n=13) and mice after 4 days of MD (P32 4d MD, n=7) (Welch’s t test).

Histograms of the distribution of ODI values for neurons from non-deprived mice at both P28 and P32 reveal the contralateral bias of neuronal responses in V1 (Figure 1D). A majority of neurons responsive to the contralateral eye did not display a significant response when the visual stimuli were presented to the ipsilateral eye, resulting in an ODI score of 1. By comparison, neurons from mice receiving 4 days of MD initiated at P28 displayed significant shifts in OD histograms driven in part by an increased percentage of ipsilateral monocular neurons with an ODI score of −1 (Figure 1D).

Experiments employing VEPs or optical imaging of intrinsic signals report that depression of responsiveness to the closed contralateral eye is the principal mechanism of OD plasticity during the critical period (Frenkel and Bear, 2004; Sato and Stryker, 2008). To probe how MD alters responsiveness at cellular resolution, we compared the dF/F responses for visual stimuli presented to either the contralateral or ipsilateral eye for non-deprived mice and those receiving 4-day MD. Consistent with these preceding measurements of aggregate neuronal activity, deprivation reduced the average response amplitude for the contralateral eye per mouse (P = .0007) but did not affect responses to the fellow ipsilateral eye (P=.79) (Figure 1E,F). This reduction of responses for the contralateral eye was evident across a range of response strengths (Figure 1G).

MD did not alter the distribution of preferred orientation but reduced binocular matching of preferred orientation (Figure 2A and S2). The matching of orientation preference of P28-P32 binocular neurons for non-deprived mice was significantly greater than P32 mice following MD (P=.018; K-S test) (Figure 2A). These results confirm findings with single-unit recordings (Wang et al., 2010). The distribution of preferred SFs for neurons were similar for the contralateral and ipsilateral eye between non-deprived mice and following MD (Figure 2B). The average preferred SF per mouse was also similar for neurons responsive to the contralateral and ipsilateral eye for both P28-P32 non-deprived mice and mice receiving 4-day MD (Figure 2B). However, despite normal distribution of preferred SF, the percentage of visually-responsive neurons for the contralateral eye that displayed significant responses for each SF was markedly reduced in P32 MD mice (P< .0002, 2-way ANOVA) (Figure 2C). This reduction of responsiveness in the population of neurons correlates with the lower range VEPs measured after MD and may contribute to lower acuity measured with behavioral assays as well (Frenkel and Bear, 2004).

**Figure 2.**
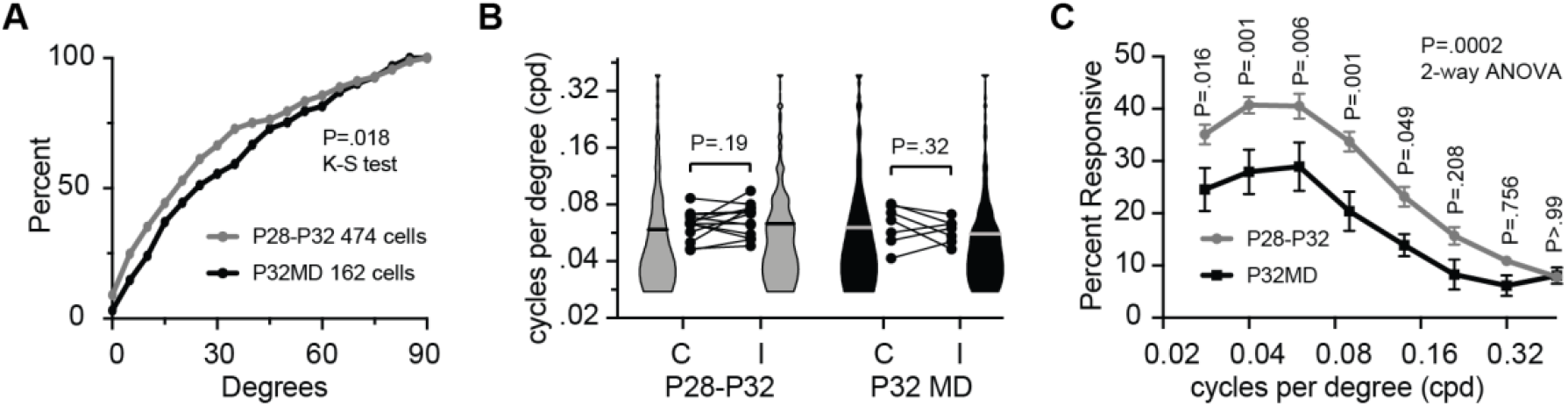
MD during the critical period reduces binocular orientation matching and the number of responding neurons across most spatial frequencies. **(A)** Cumulative distribution of difference in preferred orientation for the two eyes by binocular neurons for non-deprived mice (n= 474, median = 20 degrees) and after 4-days of MD (n=162, median = 26 degrees) (P=.018; Kolmogorov-Smirnov test of cumulative distribution). **(B)** The mean preferred SF for neurons responding to the contralateral eye (C) and ipsilateral eye (I) for each non-deprived mouse (P28-32, n=13) and mice after 4 days of MD (P32 MD, n=7). The distribution and median for the population of neurons are presented as violin plots with the median indicated by a horizontal bar. (P28-32, C = 1566 neurons, I = 766 neurons; P32MD, C = 447 neurons, I = 579 neurons). There is no statistical difference in the mean preferred SF for the C and I eye per mouse for either non-deprived mice or following 4 days of MD (paired t-test). **(C)** The percent of visually-responsive neurons per mouse that displayed a significant response to each SF at any orientation (2-way ANOVA with Sidak’s multiple comparison test for 8 comparisons).

To determine how MD during the critical period alters tuning properties of individual neurons, we measured the responses of the same neurons in L2/3 on P28 and P32 after MD (Figure 3, Figure S3). We determined the location of the same population of neurons for repeated calcium imaging by depth of focus combined with neurons with high baseline fluorescence and features of the microvasculature as reference points (Marks and Goard, 2021; Ranson, 2017; Tan et al., 2020). Given the size of neuronal soma, the magnitude of the dF/F calcium signal is resistant to minor differences in focal depth (Tan et al., 2020). However, to account for potential differences in the angle of the focal plane associated with repeatedly positioning the mouse for imaging, and to provide additional certitude that the same population of neurons was imaged in both sessions, we confined our analysis to the imaging plane circumscribed by a perimeter of neurons with tuning properties that did not change between imaging sessions (Figure S3). These neurons possessed preferred orientation that varied by less than 30 degrees (median 4 degrees, mean 7 degrees) and preferred SF that deviated by less than an octave from P28 to P32 (median .23 octaves, mean .25 octaves) (Figure S3).

**Figure 3.**
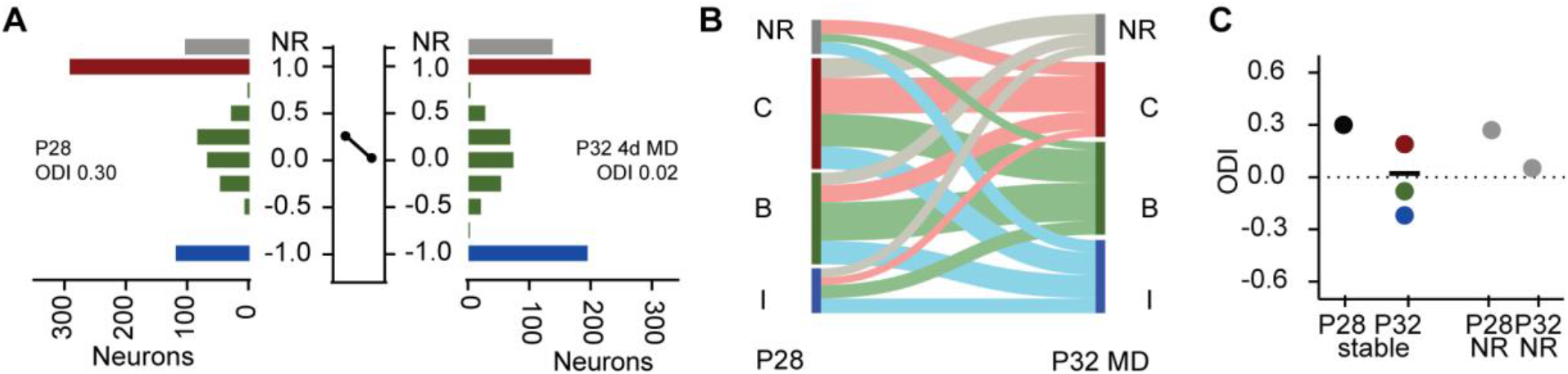
Tracking neurons longitudinally reveals an exchange of monocular and binocular neurons as well as neurons active in visual circuitry during OD plasticity. **(A)** Histogram of ODI values at P28 (*left*) and P32 4d MD (*right*) for visually-responsive neurons, non-responsive (NR) neurons at P28 that were visually-responsive at P32, and visually-responsive neurons that became non-responsive (NR) at P32, for 6 mice receiving 4 days of MD starting at P28. A line between the two histograms connects points that indicate the mean ODI of the population of visually-responsive neurons at P28 and P32MD, respectively. **(B)** Sankey diagram of the stability and interconversion between P28 and P32 MD for neurons that were non-responsive (NR, grey), monocular contralateral (C, red), binocular (B, green) and monocular ipsilateral (I, blue) for the neurons presented in panels A (**C**) The mean ODI for all neurons by category presented in panels A and D. The mean ODI of neurons that were visually-responsive at P28 (black) and the mean of ODI at P32 of these neurons categorized as monocular contralateral (red), binocular (green) and monocular ipsilateral (blue) at P28. The black horizonal line indicates the mean ODI of all neurons visually-responsive at P32 that were also visually-responsive at P28. In addition, the mean ODI of neurons at P28 which were non-responsive (NR) at P32, and the mean ODI of neurons at P32 which were non-responsive at P28 are plotted to the right.

Neurons imaged at P28 displayed significant alterations to binocularity at P32 after 4 days of MD. The distribution of ODI values shifted towards the non-deprived eye (P28, 656 neurons, mean ODI 0.30; P32, 639 neurons, mean ODI 0.02) (Figure 3A). Consistent with the OD histograms for mice imaged after MD (Figure 1D), OD plasticity decreased the ratio of monocular contralateral neurons to monocular ipsilateral neurons. A Sankey plot reveals that interconversion of neurons between monocular contralateral (294 neurons), binocular (243 neurons), and monocular ipsilateral (119 neurons) at P28 decreased the number of monocular contralateral neurons (198 neurons) and increased the number of monocular ipsilateral neurons (194 neurons) (Figure 3B). Nearly half of monocular contralateral neurons at P28 gained responsiveness to the non-deprived eye following MD. The average ODI of the population of neurons that were either monocular contralateral, binocular, and monocular ipsilateral at P28 shifted to a distribution of ODI values near zero after 4 days of MD (Figure 3C). In addition, neurons that were non-responsive at P28 but visually-responsive at P32 displayed lower average ODI values (ODI=.05) than neurons visually-responsive at P28 but non-responsive at P32 (ODI=.27). Thus, OD plasticity both converts monocular contralateral neurons into binocular neurons and monocular ipsilateral neurons, as well as recruits into visual circuitry non-responsive neurons that adopt a similar distribution of ODI values (Figure 3B,C).

OD plasticity also degraded binocular matching of preferred orientation for neurons that were binocular both at P28 (median 19 degrees) and P32 after MD (median 33 degrees) (Figure 4A). Neurons that were visually-responsive at P28 but non-responsive after MD displayed better matching of preferred orientation (median 20 degrees) than neurons non-responsive at P28 but visually-responsive at P32 (median 31 degrees) (Figure 4B). However, due to the smaller sample size (33 and 21 neurons, respectively) this comparison did not reach statistical significance (P=.29), although the medians for the two groups were similar to the total populations of binocular neurons at the two time points (P28, 18 degrees, P32MD 34 degrees) (Figure 4C).

**Figure 4.**
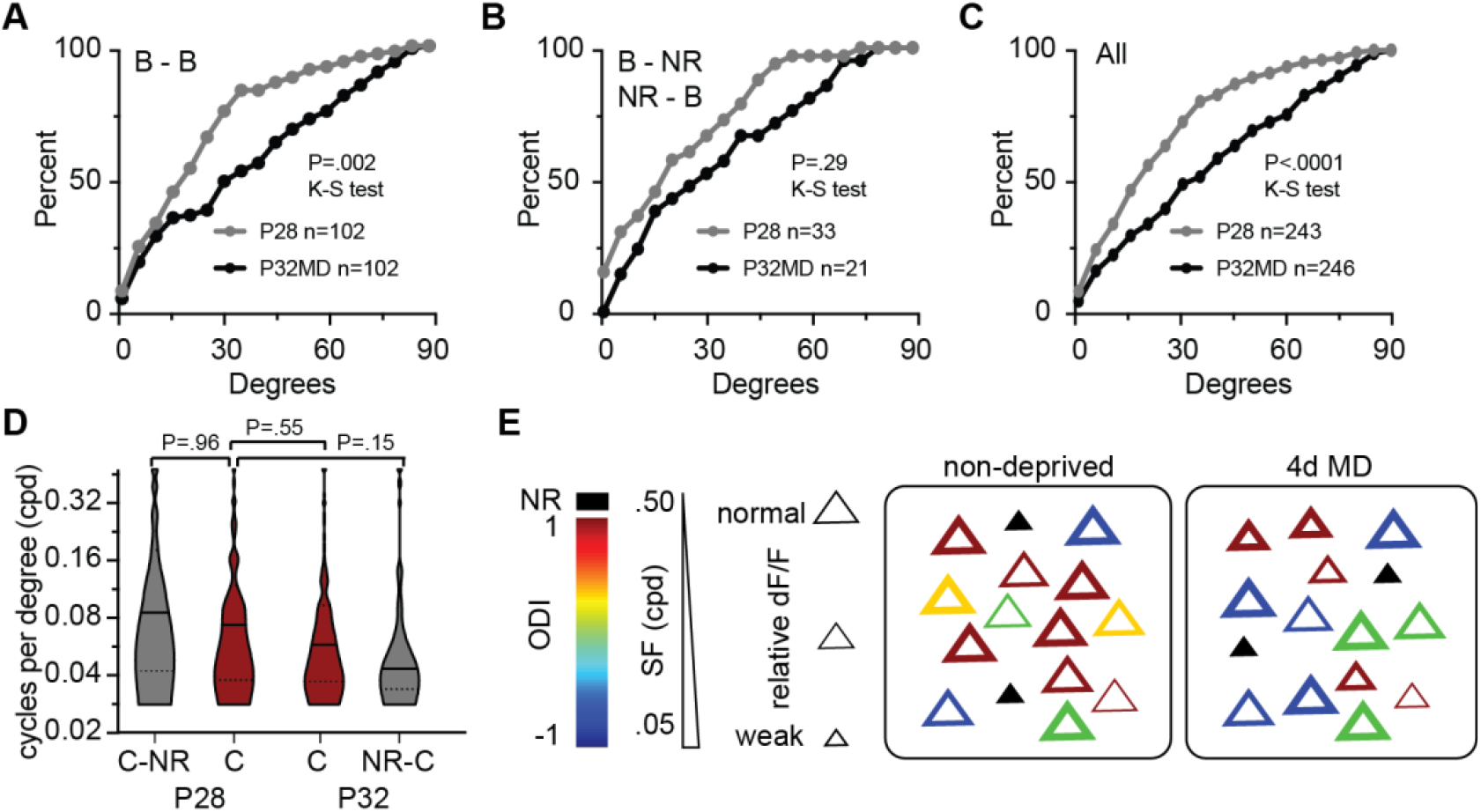
MD during the critical period degrades binocular orientation matching but does not alter the distribution of preferred SF. **(A)** Cumulative distribution of difference in preferred orientation for the two eyes by neurons that were binocular (B - B) both at P28 (n= 102, median = 19 degrees) and at P32 after 4 days of MD (n= 102, median = 34 degrees). (P<.0001; Kolmogorov-Smirnov test of cumulative distribution). (**B**) Cumulative distribution of difference in preferred orientation for the two eyes by neurons that were binocular at P28 but non-responsive at P32 (B-NR, n= 33, median = 19 degrees) and at P32 after 4 days of MD (NR-B, n= 102, median = 34 degrees). (P<.0001; Kolmogorov-Smirnov test of cumulative distribution). **(C)** Cumulative distribution of difference in preferred orientation for the two eyes by all binocular neurons (All) at P28 (n= 243, median = 19 degrees) and P32 after 4 days of MD (n= 246, median = 34 degrees). (P<.0001; Kolmogorov-Smirnov test of cumulative distribution). (**D**) The distribution and median for the population of monocular contralateral neurons presented as violin plots with the median indicated by a horizontal bar for neurons visually-responsive at P28 but non-responsive at P32 (C-NR, n= 51), neurons visually-responsive at P28 and P32 (C, n=95), neurons non-responsive at P28 and visually-responsive at P32 (NR-C, n=37). There is no statistical difference in the mean preferred SF between P28 C and the other three groups (Kruskal-Wallis test with Dunn’s correction). (**E**) Summary of alterations to neuronal tuning caused by MD during the critical period. The distributions of neuronal tuning properties are schematized for non-deprived mice and after 4 days of MD. Excitatory neurons are represented as triangles. Color corresponds to ODI. Non-responsive (NR) neurons are filled black. Preferred SF is denoted by the thickness of the colored line. Size of the triangle represents the response strength after MD relative to non-deprived.

To explore the mechanism for reduced matching of preferred orientation we compared the difference in orientation preference to the stability of preferred orientation for the contralateral and ipsilateral eye for binocular neurons (Figure S4A-C). These comparisons did not reveal any evident correlation between the precision of binocular matching of orientation preference and the stability orientation preference for either the contralateral or ipsilateral eye. In addition, we performed a similar comparison for the binocular matching of neurons that converted from monocular contralateral or monocular ipsilateral at P28 to binocular after 4 days of MD. These comparisons also did not indicate an evident correlation between binocular matching and stability of orientation preference was similar for contralateral monocular neurons and ipsilateral monocular neurons that converted to binocular (Figure S4D-F). Likewise, the stability of orientation preference for binocular neurons that converted to monocular was similar to neurons that remained monocular after MD (Figure S4G,H).

A schematic illustrates the alterations to the tuning properties and composition of visually responsive neurons following OD plasticity (Figure 4E). Monocular deprivation drives reduces the ratio of contralateral monocular neurons to ipsilateral monocular neurons with no evident change in the percentage of binocular neurons or their relative eye dominance. The relative strength of responses to the contralateral eye is diminished. Orientation preference is not represented because is unaffected by MD although binocular matching of orientation is reduced. Most neurons are tuned to SFs (~.05 cpd) that are well below the acuity threshold (~0.5 cpd). A fraction of responsive neurons displaying the contralateral bias typical of mouse visual cortex is exchanged for neurons more responsive to the non-deprived eye.

## Discussion

A current model for OD plasticity during the critical period is that occluding vision through one eye attenuates sensory input to lower the activity of neurons in V1 which in turn drives intracortical disinhibition that facilitates competitive synaptic plasticity to alter binocularity (Kuhlman et al., 2013). To understand how OD plasticity during the critical period operates at neuronal resolution, first we characterized the OD, response strength, orientation tuning, and SF tuning, for thousands of neurons and then we tracked how MD altered these properties for hundreds of neurons.

MD shifts OD towards the non-deprived eye by reducing both the number and strength of neuronal responses to the deprived eye while increasing the number of neurons responsive to the fellow eye but not their response strength. These findings explain the reduced amplitude of response to the deprived eye measured with VEPs and optical imaging of intrinsic signals (Frenkel and Bear, 2004; Sato and Stryker, 2008). Interestingly, the principal mechanism of OD plasticity was the change in the ratio of monocular neurons.

We observe that MD during the critical period did not affect orientation tuning but impaired matching of orientation preference for binocular neurons. These findings are consistent with a previous study measuring binocular matching of units isolated from electrophysiologic recordings (Wang et al., 2010). MD also did not alter the distribution of SF preference for neurons. However, it significantly reduced the percentage of the neurons responsive to the deprived eye across a range SFs. This appears to be a consequence of the overall depression of responses to the contralateral eye and may inform interpretation of VEPs across spatial frequency that have been employed to estimate acuity (Kang et al., 2013; Porciatti et al., 1999).

A preceding calcium imaging study examined the effects of a week of MD in adult mice on binocularity and orientation preference (Rose et al., 2016). They reported that adult OD plasticity is mediated by a shift in the relative responsiveness of a predominant population of binocular neurons towards the non-deprived eye, a reduction in the strength of response to the contralateral (closed) eye, and an increase in the strength of responses to the ipsilateral (non-deprived) eye. MD of adult mice did not affect orientation preference. By comparison, measurements of OD plasticity with either VEPs and optical imaging report that adult OD plasticity is mediated by strengthening of responses for the ipsilateral eye not depression of responses to the contralateral eye (Morishita and Hensch, 2008; Sato and Stryker, 2008; Sawtell et al., 2003). Some of the variability in neuronal binocularity evident in these longitudinal experiments was likely a consequence of the interconversion neurons between binocular and monocular for either eye (Tan et al., 2020). Changes to the composition of binocular circuitry are more prominent during the critical period but also substantial in adult mice (Tan et al., 2020).

Tracking the tuning properties of neurons before and after MD revealed that abnormal vision engages the synaptic mechanisms that interconvert binocular and monocular neurons and that exchange neurons active in visual circuitry. MD converted a fraction contralateral monocular neurons to binocular neurons and rendered a similar fraction of binocular neurons monocular and responsive to the contralateral and ipsilateral eye in equal proportions. This reorganization of visual circuitry reduced the ratio of monocular neurons responsive to the contralateral versus ipsilateral eye, and thereby decreased the contralateral bias that is characteristic of mouse visual cortex (Cang et al., 2005; Dräger, 1975; Porciatti et al., 1999). These alterations in the binocularity of visually responsive neurons were accompanied by an exchange of neurons active in visual circuitry that matched the altered binocularity of neurons that were visually-responsive at both P28 and at P32 after 4 days of MD. We propose that altering the tuning of responsive neurons, recruiting neurons with matching tuning properties, and a depression of the strength of responses to the contralateral eye are the neuronal basis for OD plasticity during the critical period.

## METHODS

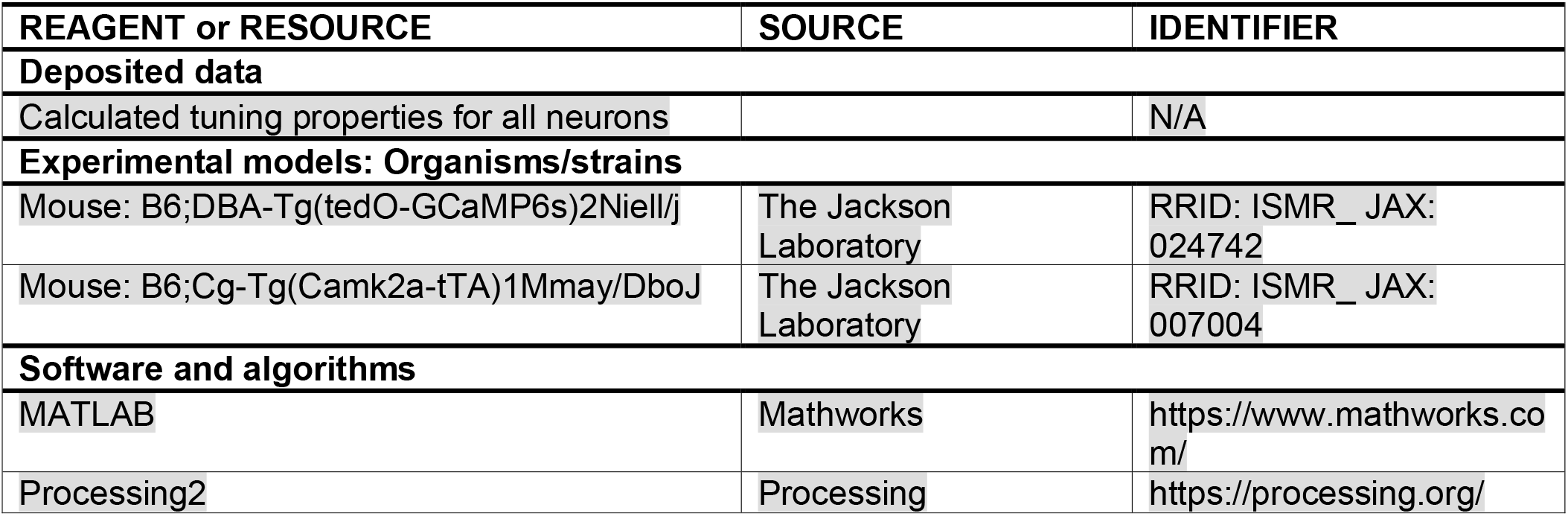

### Lead contact

Further information and requests for resources and reagents should be directed to and will be fulfilled by the Lead Contact, Aaron McGee.

### Materials availability

This study did not generate new unique reagents

### Data and Code availability

The heat maps of neuronal responses and calculated tuning properties set for this study will be deposited in Mendeley data.

## EXPERIMENTAL MODEL AND SUBJECT DETAILS

All procedures were approved by University of Louisville Institutional Animal Care and Use Committee (IACUC) and were in accord with guidelines set by the US National Institutes of Health. Mice were housed in groups of 4-5 per cage in a 12/12 light dark cycle. Animals were naive subjects with no prior history of participation in research studies. A total of 43 mice, both male (23) and female (20) were used in this study. The following male and female mice are represented in the following groups: P28-P32 non-deprived mice, 8 males and 9 females; P32 4MD, 4 males and 4 females; P28-P32 4MD repeat imaging, 5 males and 3 females; P60 non-deprived, 3 males and 2 females; P60 4MD, 2 males and 3 females.

### Mice

Imaging was performed on mice expressing GCaMP6S in excitatory neurons in forebrain. The *CaMKII-tTA* (stock no. 007004) and *TRE-GCaMP6s* (stock no. 024742) transgenic mouse lines were obtained from Jackson Labs(Mayford et al., 1996; Wekselblatt et al., 2016). Mice were genotyped with primer sets suggested by Jackson labs.

## METHODS

### Cranial window surgeries

All epifluorescent and two-photon imaging experiments were performed though a cranial window. In brief, mice were administered carprofen (5mg/kg) and buprenorphrine (0.1 mg/kg) for analgesia and anesthetized with isoflurane (4% induction, 1-2% maintenance). The scalp was shaved and mice were mounted on a stereotaxic frame with palate bar and their body temperature maintained at 37° C with a heat pad controlled by feedback from a rectal thermometer (Physitemp). The scalp was resected, the connective tissue removed from the skull, and an aluminum headbar affixed with C&B metabond (Parkell). A circular region of bone 3mm in diameter centered over left visual cortex was removed using a high-speed drill (Foredom). Care was taken to not perturb the dura. A sterile 3mm circular glass coverslip was sealed to the surrounding skull with cyanoacrylate (Pacer technology) and dental acrylic (ortho-jet, Lang Dental). The remaining exposed skull likewise sealed with cyanoacrylate and dental acrylic. Mice recovered on a heating pad. Mice were left to recover for at least 2 days prior to imaging.

### Wide-field calcium imaging

After implantation of the cranial window and before 2-photon imaging, the binocular zone of visual cortex was identified with wide-field calcium imaging similar to our method for optical imaging of intrinsic signals(Frantz et al., 2016). In brief, mice were anesthetized with isoflurane (4% induction), provided a low dose of the sedative chlorprothixene (0.5mg/kg i.p.; C1761, Sigma) and secured by the aluminum headbar. The eyes were lubricated with a thin layer of ophthalmic ointment (Puralube, Dechra Pharmaceuticals). Body temperature was maintained at 37°C with heating pad regulated by a rectal thermometer (TCAT-2LV, Physitemp). Visual stimulus was provided through custom-written software (MATLAB, Mathworks). A monitor was placed 25cm directly in front of the animal and subtended +40 to −40 degrees of visual space in the vertical axis. A horizonal white bar (2 degrees high and 20 degrees wide) centered on the zero-degree azimuth drifted from the top to bottom of the monitor with a period of 8s. The stimulus was repeated 60 times. Cortex was illuminated with blue light (475 ± 30nm) (475/35, Semrock) from a stable light source (intralux dc-1100, Volpi). Fluorescence was captured utilizing a green filter (HQ620/20) attached to a tandem lens (50mm lens, computar) and camera (Manta G-1236B, Allied Vision). The imaging plane was defocused to approximately 200 microns below the pia. Images were captured at 10Hz as images of 1024×1024 pixels and 12-bit depth. Images were binned spatially 4×4 before the magnitude of the response at the stimulus frequency (.125 Hz) was measured by Fourier analysis.

### Visual stimulus and two-photon calcium imaging

Visual stimulus presentation and image acquisition were both performed according to published methods with minor modifications (Jimenez et al., 2018; Tan et al., 2020). In brief, a battery of static sinusoidal gratings were generated in real time with custom software (Processing, MATLAB). Stimulus presentation was synchronized to the imaging data by time stamping the presentation of each visual stimulus to the image acquisition frame number a transistor-transistor logic (TTL) pulse generated with an Arduino at each stimulus transition. Orientation was sampled at equal intervals of 30 degrees from 0 to 150 degrees (6 orientations). Spatial frequency was sampled in 8 steps on a logarithmic scale at half-octaves from 0.028 to 0.48 cycles per degree. An isoluminant grey screen was included (blank) was provided as a 9^th^ step in the spatial frequency sampling as a control. Spatial phase was equally sampled at 45-degree intervals from 0 to 315 degrees for each combination of orientation and spatial frequency. Gratings with random combinations of orientation, spatial frequency, and spatial phase were presented at a rate of 4 Hz on a monitor with a refresh rate of 60Hz. Imaging sessions were 10 minutes (2400 presentations in total). Consequently, each combination of orientation and spatial frequency was presented 40 times on average (range 29-56). The monitor was centered on the zero azimuth and elevation 35cm away from the mouse and subtended 45 (vertical) by 80 degrees (horizontal) of visual space.

Imaging was performed with a resonant scanning two-photon microscope controlled by Scanbox image acquisition and analysis software (Neurolabware). The objective lens was fixed at vertical for all experiments. Fluorescence excitation was provided by a tunable wavelength infrared laser (Ultra II, Coherent) at 920 nm. Images were collected through a 16x water-immersion objected (Nikon, 0.8 NA). Images (512×796 pixels, 520×740 microns) were captured at 15.5 Hz at depths between 150 – 400 microns. Eye movements and changes in pupil size were recorded using a Dalsa Genie M1280 camera (Teledyne Dalsa) fitted with 50 mm 1.8 lens (Computar) and a 800nm long-pass filter (Edmunds Optics). Imaging was performed on alert mice positioned on a spherical treadmill by the aluminum head bar affixed to the skull. The visual stimulus was presented to each eye separately by covering the fellow eye with a small custom occluder.

### Image Processing

Image processing was performed as described previously with minor modifications (Tan et al., 2020). In summary, imaging series for each eye were motion corrected with the SbxAlign tool. Regions of interest (ROIs) corresponding to excitatory neurons were selected manually with the SbxSegment tool following computation of pixel-wise correlation of fluorescence changes over time from 350 evenly spaced frames (~4%). ROIs for each experiment were determined by correlated pixels the size similar to that of a neuronal soma. The fluorescence signal for each ROI and the surrounding neuropil were extracted from this segmentation map.

### Image analysis to identify visually responsive neurons and calculate their tuning properties

Image analysis was performed as described previously with minor modifications (Tan et al., 2020). The fluorescence signal for each neuron was extracted by computing the mean of the calcium fluorescence within each ROI and subtracting the median fluorescence from the surrounding perimeter of neuropil (Ringach et al., 2016; Tan et al., 2020). An inferred spike rate (ISR) was estimated from adjusted fluorescence signal with the Vanilla algorithm (Berens et al., 2018). A reverse correlation of the ISR to stimulus onset was used to calculate the preferred stimuli (Jimenez et al., 2018; Ringach et al., 2016; Tan et al., 2020; Yaeger et al., 2019). Neurons that satisfied three criteria were categorized as visually responsive: (1) the ISR was highest with the optimal delay of 4-9 frames following stimulus onset. This delay was determined empirically for this transgenic GCaMP6s mouse (Tan et al., 2020). (2) the SNR was at least one standard deviation greater than spontaneously active neurons. The signal is the mean of the spiking standard deviation at the optical delay between 4-9 frames after stimulus onset and the noise this value at frames −2 to 0 before the stimulus onset or 15-18 after it (Jimenez et al., 2018; Tan et al., 2020) (3) and the percent of responses to the preferred stimulus was at least one standard deviation greater than spontaneously active neurons (See Suppl. Fig 1 k-n). The visual stimulus capturing the preferred orientation and SF was the determined from the matrix of all orientations and SFs presented as the combination with highest average ISR.

The preferred orientation for each neuron was calculated as:

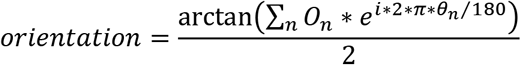

On is a 1×6 array of the mean z-scores associated with the calculation of the ISR at orientations Q_n_ (0 to 150 degrees, spaced every 30 degrees). Orientation calculated with this formula is in radians and was converted to degrees. The tuning width was the full width at half-maximum of the preferred orientation. The preferred SF for each neuron was calculated as:

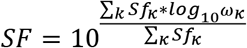

Sf_k_ is a 1×8 array of the mean z-scores at SFs w_k_ (8 equal steps on a logarithmic scale from 0.028 to 0.481 cycles per degree). Tails of the distribution were clipped at 25% of the peak response. The tuning width was the full width at half-maximum of the preferred SF in octaves. The percent visually responsive neurons with significant responses at each SF was determined by comparing the distribution of ISR values at each SF versus the stimulus blank with a KW-test with Dunn’s correction for 8 comparisons. Neurons with P<.01 for a given SF were considered significance responses at that SF.(Salinas et al., 2017)

### Ocular Dominance Index (ODI)

ODI was calculated as (C−I)/(C+I), where C and I are the mean normalized change in fluorescence (dF/F) for the preferred visual stimulus for the contralateral eye and ipsilateral eye, respectively. In cases where neurons displayed no significant response to visual stimuli provided to one eye, they were considered monocular for the other eye and assigned ODI values of 1 (contralateral) and −1 (ipsilateral)(Salinas et al., 2017).

### Longitudinal imaging of neurons before and after MD

The same imaging plane was identified for the second imaging experiment by using the reference image of the first experiment. The location in the neuropil was achieved predominantly by coordinating the depth from pial surface and the position of small blood vessels (Suppl. Fig. 3a). Fine adjustment of position was performed by matching responsive neurons evident in the reference image from the first experiment while presenting the visual stimulus.

Each imaging session was segmented independently and every ROI was assigned a unique number. No geometric transformations were performed to match segmentation masks for ROIs from the two imaging sessions. The segmentation masks for the two imaging sessions were then compared and ROIs with at least 50% overlap were considered the same neuron. A perimeter of neurons with overlapping ROIs and tuning properties that did not change between imaging sessions (a difference in orientation preference of less than 30 degrees and SF preference of less than a half octave) defined the matched imaging plane. To determine the SNR values of lost neurons at P32 and gained neurons at P28, the segmentation masks were exchanged between time points and the SNR from ROIs for the corresponding neurons at the other time point were calculated.

### Monocular deprivation

The right eye was sutured closed with a single mattress suture with 6-0 Prolene monofilament (Ethicon 8709) (Stephany et al., 2014). Prior to imaging, mice were briefly (< 5 min) anesthetized with isoflurane (4% induction, 1-2% maintenance), the suture removed with Vannas scissors (Fine Science Tools). The eye was flushed with sterile saline and examined for corneal abrasions with a stereomicroscope. The mouse was then immediately head-fixed for imaging and allowed to recover for no less than 45 minutes. The occluder was positioned over one eye as soon as the mouse was head-fixed and occluded one of the eyes at all times. At no point during the experiment were mice permitted unobstructed binocular vision.

### Statistics

No statistical methods were used to predetermine sample size. All statistical analyses were done using Prism 8 software (GraphPad Software). All data sets were examined for normality and two-tail t-tests were only employed for data with normal distributions. The non-parametric Mann-Whitney (MW) test for pairwise comparisons and Kruskal-Wallis (KW) test with Dunn’s correction for multiple comparisons were employed for data that did not conform to a normal distribution.

## Acknowledgements

We thank D. Ringach and J. Trachtenberg for sharing software and hardware design for visual stimulus presentation and data analysis prior to publication, A. Eliasen and G. Armstrong for software development, and B. Croslin for mouse husbandry and genotyping. This research is supported by the National Eye Institute (R01EY021580 and R01EY027407).

## Author Contributions

The contributions are as follows: TCB and AWM designed the study. TCB performed the experiments. TCB and AWM analyzed the data, built the figures, and wrote the manuscript.

## Declaration of Interests

The authors declare no competing interests

## Data and Materials Availability

Requests should be addressed to the corresponding author A.W. McGee.

## Notes

### Competing Interest Statement

The authors have declared no competing interest.

## References

Berens P, Freeman J, Deneux T, Chenkov N, McColgan T, Speiser A, Macke JH, Turaga SC, Mineault P, Rupprecht P, Gerhard S, Friedrich RW, Friedrich J, Paninski L, Pachitariu M, Harris KD, Bolte B, Machado TA, Ringach D, Stone J, Rogerson LE, Sofroniew NJ, Reimer J, Froudarakis E, Euler T, Román Rosón M, Theis L, Tolias AS, Bethge M. 2018. Community-based benchmarking improves spike rate inference from two-photon calcium imaging data. PLoS Comput Biol 14:1–13. doi:10.1371/journal.pcbi.1006157

Cang J, Kalatsky VA, Löwel S, Stryker MP. 2005. Optical imaging of the intrinsic signal as a measure of cortical plasticity in the mouse. Vis Neurosci 22:685–691. doi:10.1017/S0952523805225178

Crair MC, Ruthazer ES, Gillespie DC, Stryker MP. 1997. Relationship between the ocular dominance and orientation maps in visual cortex of monocularly deprived cats. Neuron 19:307–318. doi:10.1016/S0896-6273(00)80941-1

Drager UC. 1978. Observations on monocular deprivation in mice. J Neurophysiol 41:28–42. doi:10.1152/jn.1978.41.1.28

Dräger UC. 1975. Receptive fields of single cells and topography in mouse visual cortex. J Comp Neurol 160:269–289. doi:10.1002/cne.901600302

Durand S, Iyer R, Mizuseki K, De Vries S, Mihalas S, Reid RC. 2016. A comparison of visual response properties in the lateral geniculate nucleus and primary visual cortex of awake and anesthetized mice. J Neurosci 36:12144–12156. doi:10.1523/JNEUROSCI.1741-16.2016

Espinosa JS, Stryker MP. 2012. Development and Plasticity of the Primary Visual Cortex. Neuron 75:230–249. doi:10.1016/j.neuron.2012.06.009

Fagiolini M, Pizzorusso T, Berardi N, Domenici L, Maffei L. 1994. Functional postnatal development of the rat primary visual cortex and the role of visual experience: Dark rearing and monocular deprivation. Vision Res 34:709–720. doi:10.1016/0042-6989(94)90210-0

Frantz MG, Kast RJ, Dorton HM, Chapman KS, McGee AW. 2016. Nogo Receptor 1 Limits Ocular Dominance Plasticity but not Turnover of Axonal Boutons in a Model of Amblyopia. Cereb Cortex 26. doi:10.1093/cercor/bhv014

Frenkel MY, Bear MF. 2004. How monocular deprivation shifts ocular dominance in visual cortex of young mice. Neuron 44:917–923. doi:10.1016/j.neuron.2004.12.003

Gordon JA, Stryker MP. 1996. Experience-dependent plasticity of binocular responses in the primary visual cortex of the mouse. J Neurosci 16:3274–3286. doi:10.1523/jneurosci.16-10-03274.1996

Hensch TK, Fagiolini M, Mataga N, Stryker MP, Baekkeskov S, Kash SF. 1998. Local GABA circuit control of experience-dependent plasticity in developing visual cortex. Science (80-) 282:1504–1508. doi:10.1126/science.282.5393.1504

Hensch TK, Quinlan EM. 2018. Critical periods in amblyopia. Vis Neurosci 35:E014. doi:10.1017/S0952523817000219

Hubel DH, Wiesel TN. 1970. The period of susceptibility to the physiological effects of unilateral eye closure in kittens. J Physiol 206:419–436. doi:10.1113/jphysiol.1970.sp009022

Hubel DH, Wiesel TN, LeVay S. 1977. Plasticity of ocular dominance columns in monkey striate cortex. Philos Trans R Soc Lond B Biol Sci 278:377–409. doi:10.1098/rstb.1977.0050

Jenks KR, Shepherd JD. 2020. Experience-Dependent Development and Maintenance of Binocular Neurons in the Mouse Visual Cortex. Cell Rep 30:1982–1994.e4. doi:10.1016/j.celrep.2020.01.031

Jeon BB, Swain AD, Good JT, Chase SM, Kuhlman SJ. 2018. Feature selectivity is stable in primary visual cortex across a range of spatial frequencies. Sci Rep 8:1–14. doi:10.1038/s41598-018-33633-2

Jimenez LO, Tring E, Trachtenberg JT, Ringach DL. 2018. Local tuning biases in mouse primary visual cortex. J Neurophysiol 120:274–280. doi:10.1152/jn.00150.2018

Kang E, Durand S, LeBlanc JJ, Hensch TK, Chen C, Fagiolini M. 2013. Visual acuity development and plasticity in the absence of sensory experience. J Neurosci 33:17789–17796. doi:10.1523/JNEUROSCI.1500-13.2013

Kuhlman SJ, Olivas ND, Tring E, Ikrar T, Xu X, Trachtenberg JT. 2013. A disinhibitory microcircuit initiates critical-period plasticity in the visual cortex. Nature 501:543–546. doi:10.1038/nature12485

Levelt CN, Hübener M. 2012. Critical-Period Plasticity in the Visual Cortex. Annu Rev Neurosci 35:309–330. doi:10.1146/annurev-neuro-061010-113813

Marks TD, Goard MJ. 2021. Stimulus-dependent representational drift in primary visual cortex. Nat Commun 12:1–16. doi:10.1038/s41467-021-25436-3

Marshel JH, Garrett ME, Nauhaus I, Callaway EM. 2011. Functional Specialization of Seven Mouse Visual Cortical Areas. Neuron 72:1040–1054. doi:10.1016/j.neuron.2011.12.004

Mayford M, Bach ME, Huang YY, Wang L, Hawkins RD, Kandel ER. 1996. Control of memory formation through regulated expression of a CaMKII transgene. Science (80-) 274:1678–1683. doi:10.1126/science.274.5293.1678

Morishita H, Hensch TK. 2008. Critical period revisited: impact on vision. Curr Opin Neurobiol 18:101–107. doi:10.1016/j.conb.2008.05.009

Mrsic-Flogel TD, Hofer SB, Ohki K, Reid RC, Bonhoeffer T, Hübener M. 2007. Homeostatic Regulation of Eye-Specific Responses in Visual Cortex during Ocular Dominance Plasticity. Neuron 54:961–972. doi:10.1016/j.neuron.2007.05.028

Niell CM, Stryker MP. 2008. Highly selective receptive fields in mouse visual cortex. J Neurosci 28:7520–7536. doi:10.1523/JNEUROSCI.0623-08.2008

Pham TA, Graham SJ, Suzuki S, Barco A, Kandel ER, Gordon B, Lickey ME. 2004. A semi-persistent adult ocular dominance plasticity in visual cortex is stabilized by activated CREB. Learn Mem 11:738–747. doi:10.1101/lm.75304

Porciatti V, Pizzorusso T, Maffei L. 1999. The visual physiology of the wild type mouse determined with pattern VEPs. Vision Res 39:3071–3081. doi:10.1016/S0042-6989(99)00022-X

Ranson A. 2017. Stability and Plasticity of Contextual Modulation in the Mouse Visual Cortex. Cell Rep 18:840–848. doi:10.1016/j.celrep.2016.12.080

Ringach DL, Mineault PJ, Tring E, Olivas ND, Garcia-Junco-Clemente P, Trachtenberg JT. 2016. Spatial clustering of tuning in mouse primary visual cortex. Nat Commun 7:1–9. doi:10.1038/ncomms12270

Rose T, Jaepel J, Hubener M, Bonhoeffer T. 2016. Cell-specific restoration of stimulus preference after monocular deprivation in the visual cortex. Science (80-) 352:1319–1322. doi:10.1126/science.aad3358

Salinas KJ, Velez DXF, Zeitoun JH, Kim H, Gandhi SP. 2017. Contralateral bias of high spatial frequency tuning and cardinal direction selectivity in mouse visual cortex. J Neurosci 37:10125–10138. doi:10.1523/JNEUROSCI.1484-17.2017

Sato M, Stryker MP. 2008. Distinctive features of adult ocular dominance plasticity. J Neurosci 28:10278–10286. doi:10.1523/JNEUROSCI.2451-08.2008

Sawtell NB, Frenkel MY, Philpot BD, Nakazawa K, Tonegawa S, Bear MF. 2003. NMDA receptor-dependent ocular dominance plasticity in adult visual cortex. Neuron 38:977–985. doi:10.1016/S0896-6273(03)00323-4

Scholl B, Pattadkal JJ, Priebe NJ. 2017. Binocular disparity selectivity weakened after monocular deprivation in mouse V1. J Neurosci 37:6517–6526. doi:10.1523/JNEUROSCI.1193-16.2017

Stephany C-E, Ikrar T, Nguyen C, Xu X, McGee AW. 2016. Nogo Receptor 1 Confines a Disinhibitory Microcircuit to the Critical Period in Visual Cortex. J Neurosci 36:11006–11012. doi:10.1523/JNEUROSCI.0935-16.2016

Stephany CÉ, Chan LLH, Parivash SN, Dorton HM, Piechowicz M, Qiu S, McGee AW. 2014. Plasticity of binocularity and visual acuity are differentially limited by nogo receptor. J Neurosci 34:11631–11640. doi:10.1523/JNEUROSCI.0545-14.2014

Tan L, Tring E, Ringach DL, Zipursky SL, Trachtenberg JT. 2020. Vision Changes the Cellular Composition of Binocular Circuitry during the Critical Period. Neuron 1–13. doi:10.1016/j.neuron.2020.09.022

Wang BS, Sarnaik R, Cang J. 2010. Critical Period Plasticity Matches Binocular Orientation Preference in the Visual Cortex. Neuron 65:246–256. doi:10.1016/j.neuron.2010.01.002

Wekselblatt JB, Flister ED, Piscopo DM, Niell CM. 2016. Large-scale imaging of cortical dynamics during sensory perception and behavior. J Neurophysiol 115:2852–2866. doi:10.1152/jn.01056.2015

Wiesel TN, Hubel DH. 1963. Single-Cell Responses in Striate Cortex of Kittens Deprived of Vision in One Eye. J Neurophysiol 26:1003–1017. doi:10.1152/jn.1963.26.6.1003

Yaeger CE, Ringach DL, Trachtenberg JT. 2019. Neuromodulatory control of localized dendritic spiking in critical period cortex. Nature 567:100–104. doi:10.1038/s41586-019-0963-3

