## Supplemental Figures for "Monocular deprivation during the critical period alters neuronal tuning and the composition of visual circuitry"

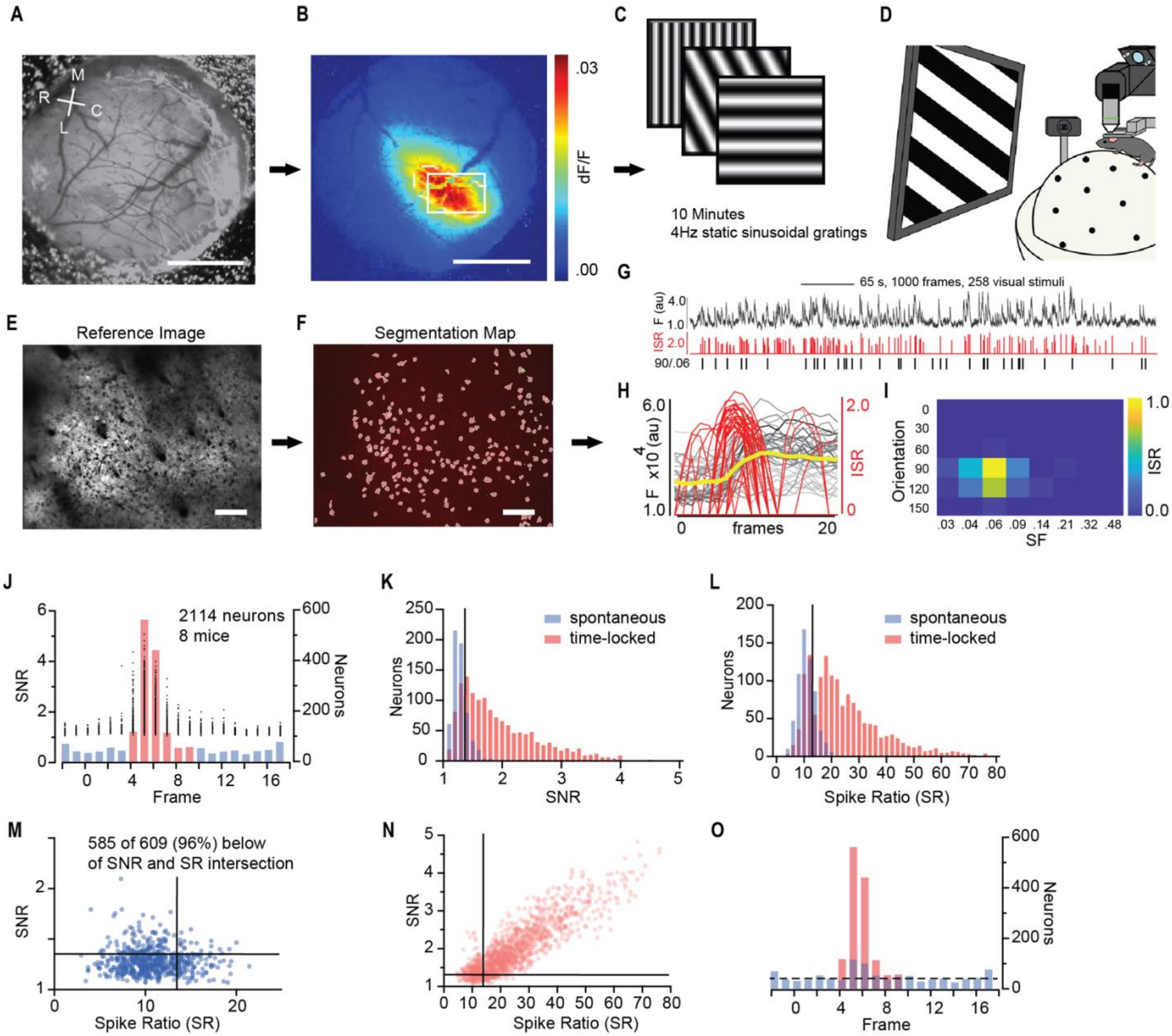

**Figure S1. Example of cranial window, identification of the binocular zone of V1, cell segmentation, signal extraction, deconvolution, and identification of visually-responsive neurons.**
**(A**) A cranial window 3 mm in diameter implanted over visual cortex. Scale bar = 1cm. Rostral – caudal and medial – lateral axes are indicated at upper left. **(B)** Wide-field calcium imaging of neuronal activity in response to a horizonal bar 30 degrees wide and 2 degrees high drifting down at 10 degrees per second. The white rectangle indicates the imaging field in (C) and (D). Scale bar = 1cm. **(C)** Schematic of the visual stimulus. Sinusoidal gratings at 30 degrees intervals in orientation and between .028 and .48 cycles per degrees in spatial frequency (SF) spaced at half octaves (log(1.5)) as well as an iso-luminant grey screen are presented in random order at 4Hz for 10 minutes. Each combination of orientation and SF is presented 40 times on average (range 29-56). **(D)** Schematic of the set up for calcium imaging of alert mice. The monitor is positioned 35 cm away from the mouse centered at the zero azimuth and elevation. A mouse is head-fixed on a styrofoam ball floating on column of air. Mouse head-fixed, alert, and freely moving. A camera records pupil diameter. **(E)** An example reference image of imaging plane in the binocular zone of visual cortex. This is the P28 image in Figure 1A. Imaging field is 750 μm X 500 μm. Scale bar = 100 μm. **(F**) Segmented neurons from the imaging field in (C). White circles correspond to ROIs identified manually. A total of 215 neuronal ROIs are segmented in this field. Scale bar = 100 μm. **(G)** Representative calcium trace (black line, *top*) and inferred spike rate (ISR) (red line, *middle*) from example neuron from above imaging field. The timing of presentations of the preferred stimulus (90 degrees, .06 cpd) during the experiment (black vertical lines, *bottom*). Images are collected at 15.5 Hz. The scale bar represents 65s, 1000 frames, and approximately 258 visual stimuli (grey horizonal bar, *top right*). Entire trace represents 10 minutes during which 2400 gratings were presented from 56 combinations (6 orientations and a grey screen for each of 8 SFs). **(H)** Fluorescent traces (grey lines) superimposed for the 20 frames (1.25 seconds) following the onset of the 36 presentations of the preferred visual stimulus for a representative neuron. Red lines represent the positions of inferred spikes. The yellow line indicates the average fluorescence across all frames and presentations. **(I)** Heat map of inferred spike rate (ISR) for all combinations of orientation and SF (in cpd) for a representative neuron. **(J)** Distribution of SNR values (black circles) for 2114 ROIs from 8 P28 non-deprived mice. SNR is plotted (*left*) versus the frame with the optimal delay. The total number of neurons with an optical delay is plotted (*right*) versus the frame number. Frames 4-9 correspond to the optimal delay of time-locked neurons (red) while frame outside this range are spontaneously active (blue). **(K)** The SNR of spontaneously active neurons with optimal delays outside of frames 3-10 (blue) and the SNR of time-locked neurons with optical delays between frames 4-9. The black vertical line indicates the threshold of the 75$^{th}$ percentile of the SNR of spontaneously active neurons. **(L)** The Spike Ratio (SR) is the percent of presentations of the optimal visual stimulus with an ISR greater than zero at the frame of optimal delay for spontaneous and time-locked neurons in (J). The black line represents the threshold of the 75$^{th}$ percentile of the SR of spontaneously active neurons. **(M)** Scatter plot of SNR versus SR for spontaneously active neurons. More than 96% are below the intersection of these criteria for visual responsiveness. **(N)** Scatter plot of SNR and percent responses for time-locked neurons. (**O)** Panel J replotted with the visually-responsive neurons (red) and spontaneously active neurons (blue) indicated. The dashed line represents the average number of spontaneously active neurons with optimal delays at outside of frames 3-10. Note that the number of spontaneously active neurons between frames 4-9 equals or exceeds that of the surrounding frames. The y-axis is shown on the right to match panel (J).

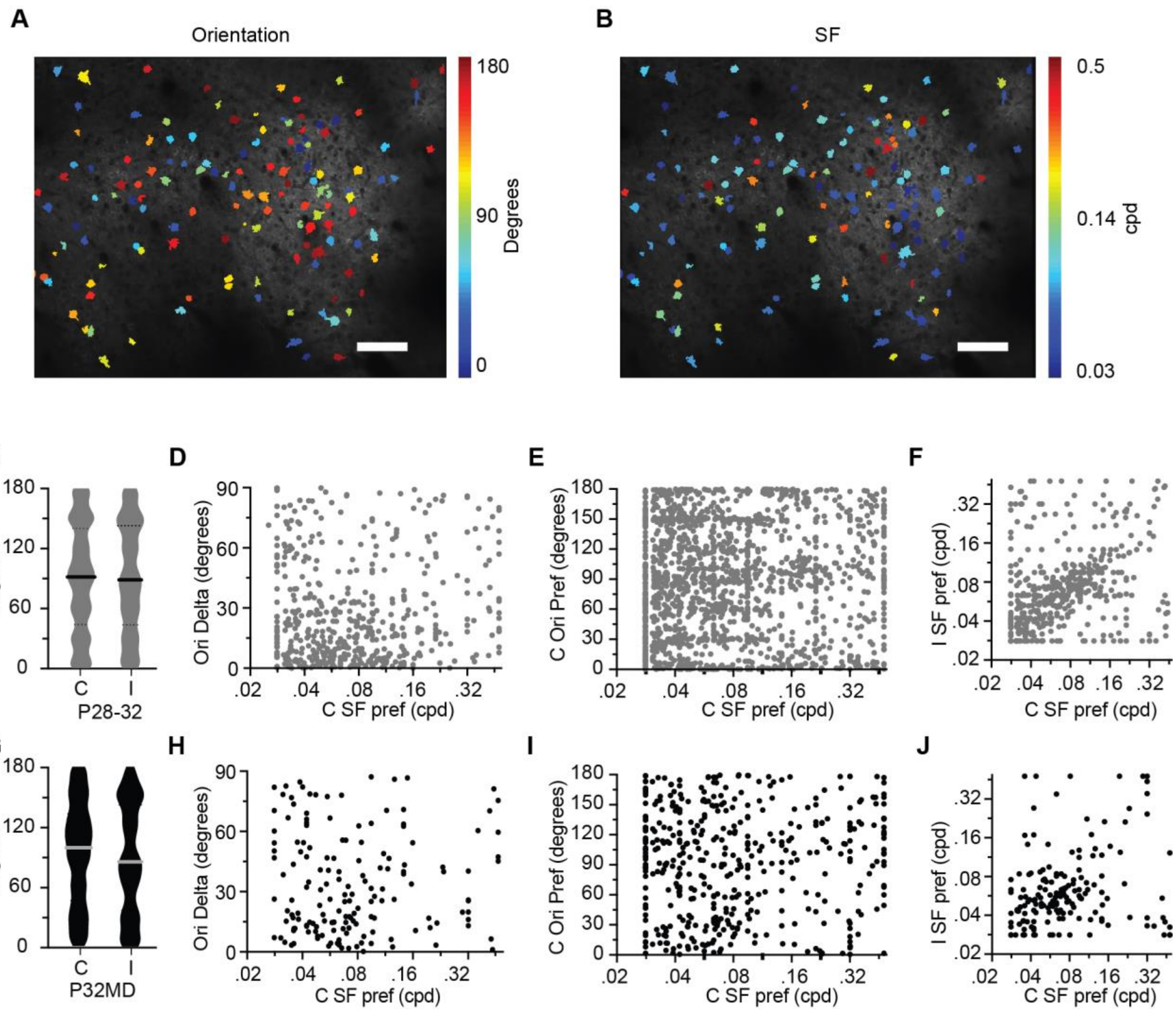


**Figure S2. The effects of MD on preferred orientation and preferred SF tuning properties of neurons for mice during the critical period**

**(A)** Heat map of neuronal orientation preference for the contralateral eye for the P28 neurons presented in Figure 1A. Scale bar equals 100μm. **(B)** Heat map of neuronal orientation preference for the contralateral eye for the P28 neurons presented in Figure 1A. Scale bar equals 100μm. **(C)** Preferred orientation for the contralateral eye (C) and ipsilateral eye (I) for all non-deprived P28-P32 mice in Figure 1A. **(D)** Difference in the preferred orientation for the contralateral eye and ipsilateral eye by binocular neurons (n=474) plotted against the preferred SF for the contralateral eye for non-deprived P28-P32 mice. **(E)** Preferred orientation plotted against preferred SF for the contralateral eye for neurons (n=1566) from non-deprived P28-P32 mice. **(F)** Preferred orientation for the ipsilateral eye plotted against preferred SF for the contralateral eye for non-deprived P28-P32 mice. **(G)** Preferred orientation for the contralateral eye (C) and ipsilateral eye (I) for P32 4-day MD mice (P32MD) in Figure 1A. **(H)** Difference in the preferred orientation for the contralateral eye and ipsilateral eye by binocular neurons (n=162) plotted against the preferred SF for the contralateral eye for P32 4-day MD mice. **(I)** Preferred orientation plotted against preferred SF for the contralateral eye for P32 4-day MD mice (n=447). **(J)** Preferred orientation for the ipsilateral eye plotted against preferred SF for the contralateral eye for for P32 4-day MD mice.

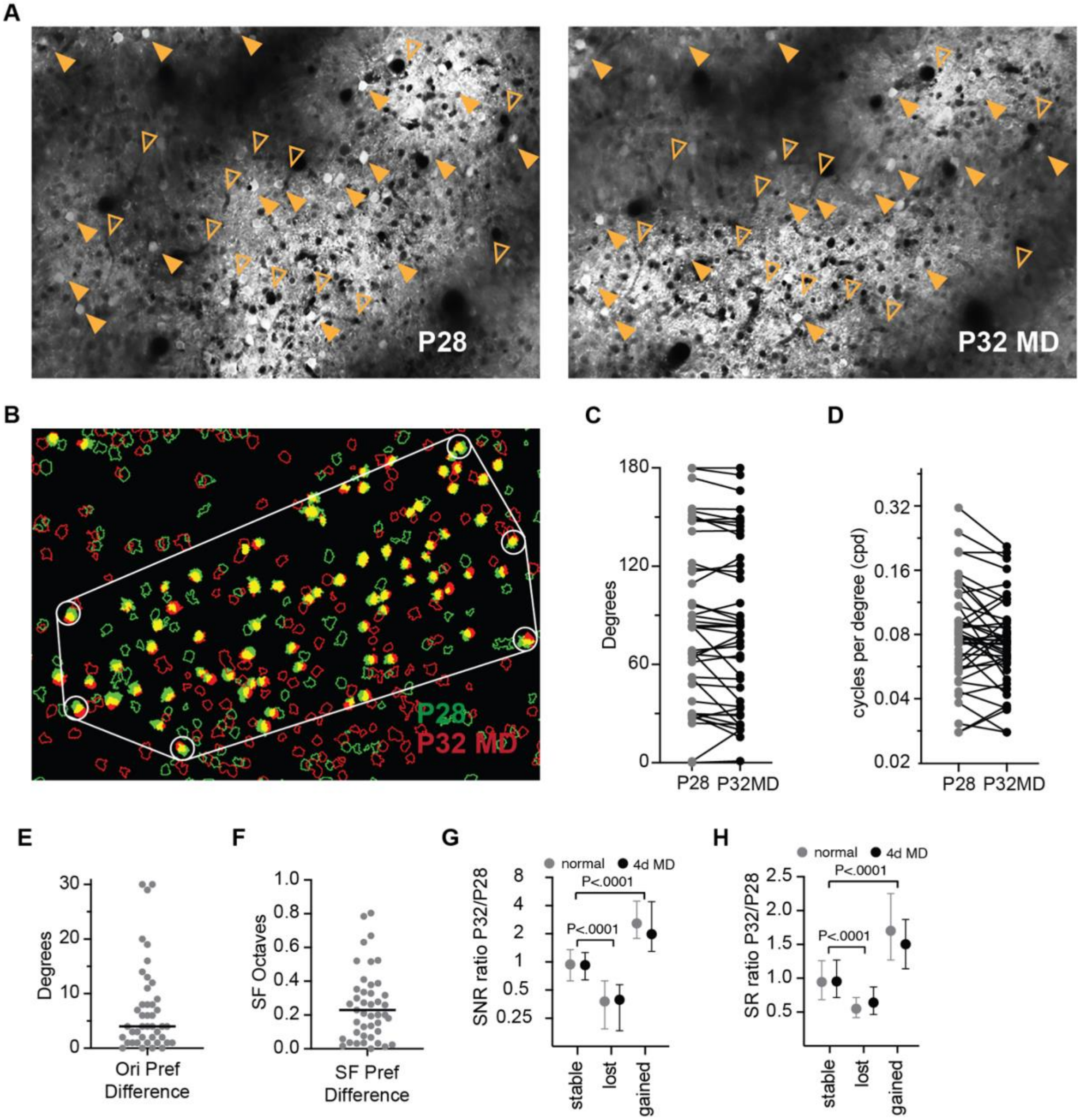


**Figure S3. Repeated calcium imaging of neurons in V1 to measure the stability of tuning properties and the neuronal composition of visual circuitry. (A)** Example reference images for the imaging plane of neurons at P28 (*right*) and P32 after 4 days of MD (*left*). Scale bar =100μm. Landmarks of strongly responding neurons (gold filled arrowheads) and features of the microvasculature (gold open arrowheads) are used to identify the same location. **(B)** Populations of segmented ROIs at P28 (green outlines) and P32 MD (red outlines). ROIs with at least 50% overlap are filled (yellow). A perimeter of overlapping ROIs subsequently determined to be visually-responsive at both time points and possess an orientation preference that differs by less than 30 degrees and SF preference that differs by less than one octave are circled (white outline). These neurons define region the analysis. **(C)** The preferred orientation of perimeter neurons at P28 and P32MD. Black lines connect pairs. **(D)** The preferred SF of perimeter neurons at P28 and P32MD. Black lines connect pairs. **(E)** Difference in the preferred orientation of perimeter neurons at P28 and P32MD. **(F)** Difference in the preferred SF of perimeter neurons at P28 and P32MD. **(G)** The P32/P28 SNR ratio of neurons which were visually-responsive at both P28 and P32 (stable), neurons that were visually-responsive at P28 but not P32, and neurons that were not visually-responsive at P28 but were visually responsive at P32. (Kruskal-Wallis Test with Dunn's correction). **(H)** The P32/P28 Spike Ratio (SR) ratio of neurons which were visually-responsive at both P28 and P32 (stable), neurons that were visually-responsive at P28 but not P32, and neurons that were not visually-responsive at P28 but were visually responsive at P32. (Kruskal-Wallis Test with Dunn's correction).

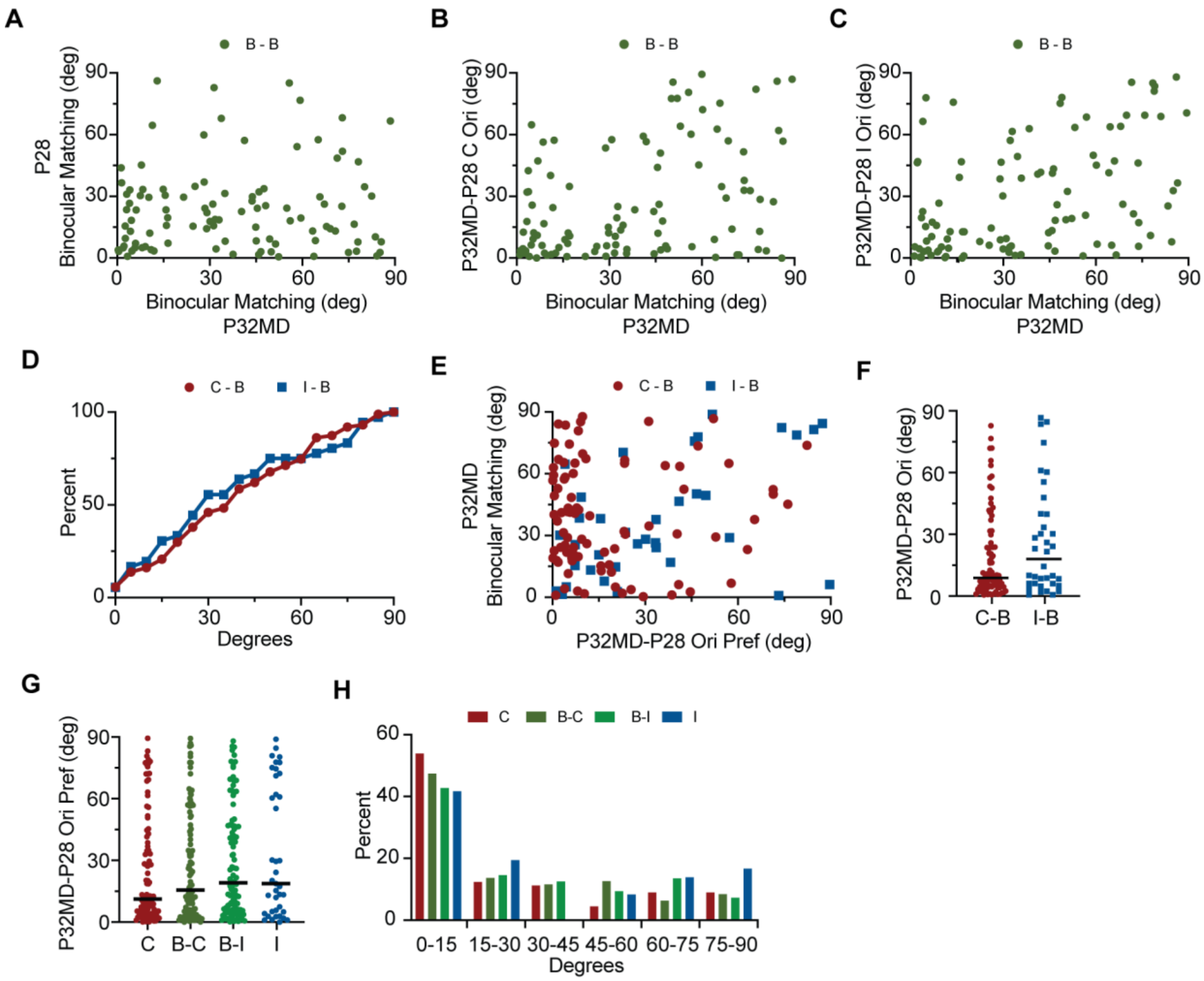


**Figure S4. Comparisons of preferred orientation, matching of preferred orientation, and preferred SF for mice receiving 4 days of MD. (A)** A scatter plot of the difference in preferred orientation (binocular matching) for neurons that were binocular both at P28 and after 4 days of MD (B-B, n=102). **(B)** The difference in preferred orientation (binocular matching) plotted against the difference in preferred orientation at P28 and P32MD for the contralateral eye. (**C)** The difference in preferred orientation (binocular matching) plotted against the difference in preferred orientation at P28 and P32MD for the ipsilateral eye. **(D)** Binocular matching difference for neurons that interconverted to binocular from monocular contralateral (red) (C-B, n=87) and monocular ipsilateral (blue) (I-B, n=36). **(E)** The difference in preferred orientation (binocular matching) from panel D plotted against the difference in preferred orientation at P28 and P32MD for P28 contralateral monocular (red) and P28 ipsilateral monocular (blue) neurons. **(F)** The difference in preferred orientation at P28 and P32MD for contralateral monocular neurons (C-B, red) and ipsilateral monocular neurons (I-B, blue) that converted to binocular neurons at P32 after 4 days of MD from panels D and E. **(G)** The difference in preferred orientation for neurons that were monocular contralateral (red, n=95), binocular (green, n=102), or monocular ipsilateral (blue, n=39) at both P28 and P32 after 4 days of MD. The preferred orientation for the contralateral eye and ipsilateral eye are shown separately for binocular neurons (B-C and B-I, respectively). **(H)** A histogram of the difference in preferred orientation presented in panel F in 15-degree intervals.
